# A droplet-to-data workflow for antimicrobial screening of complex matrices

**DOI:** 10.64898/2026.10.28.762405

**Authors:** Immanuel Sanka, Merike Vaher, Aditya Aryandi Setiawibawa, Piret Saar-Reismaa, Pille Pata, Katri Rüüson, Ott Scheler, Simona Bartkova

## Abstract

Screening plant extracts and other complex natural matrices for antibacterial activity is complicated by optical interference and population averaging in conventional bulk assays. Droplet-based assays mitigate interference while enabling high-throughput assessment of bacterial subpopulations. However, analyzing the resulting image data involves difficult image processing, classification, and interpretation. Here, we present a democratized workflow for antimicrobial screening in complex matrices that combines a polydisperse droplet-based assay with EasyFlow 2.0 (EF2), our updated droplet image analysis platform. EF2 offers stable containerized deployment, expanded polydisperse droplet support, automated thresholding and MIC estimation, LLM-assisted data exploration, and import of other software outputs such as CellProfiler™ (CP) measurements and CellProfiler Analyst™ (CPA) classification models. As proof of concept, we applied this workflow to assess the antibacterial and aggregation-modulating effects of chemically distinct extracts of *Taraxacum officinale, Rhodiola rosea*, and *Lonicera caerulea* against *Escherichia coli*. Extracts were chemically characterized by HPLC-MS and evaluated using both standard 96-well plate microdilution and droplet-based assays. In total, 27,059 droplets were segmented in CP, classified into four growth phenotypes using a CPA random forest model, and analyzed in EF2. The droplet-based assay reached minimal inhibitory concentration (MIC) at lower extract concentrations than 96-well plate microdilution and revealed extract-specific aggregation effects: *L. caerulea* reduced *E. coli* aggregation, whereas *T. officinale* did not. *R. rosea* reduced growth in the 96-well plate microdilution assay, but high concentrations caused droplet gelation that prevented droplet-based analysis, showing that sample components must be checked for compatibility with droplet formation. These results illustrate that the EF2-centered workflow supports rapid, automated MIC estimation through unsupervised thresholding when labels are unavailable, and four-class aggregation phenotyping when a CPA model is supplied. This workflow offers flexible, scalable screening, potentially extendable beyond plant extracts to other complex sample matrices.

## INTRODUCTION

Antimicrobial resistance (AMR) has become a rising global health threat.^1^ It already causes an estimated 700,000 deaths worldwide each year, a figure projected to reach 10 million by 2050.^2^ Plant extracts are gaining renewed attention for their antimicrobial properties (largely deriving from secondary metabolites such as tannins, iridoids, alkaloids, phenolic compounds, and flavonoids)^3–6^ as a promising approach to reduce antibiotic overuse.^7^

Dandelion (*Taraxacum officinale*), blue honeysuckle (*Lonicera caerulea*), and golden root (*Rhodiola rosea*) were selected as representative, locally accessible Nordic-Baltic medicinal plants with established but chemically distinct antibacterial profiles. Dandelion, an edible *Asteraceae* common in Estonian meadows, has traditional use against inflammatory and bacterial conditions, with phenolic-rich leaf and flower extracts previously shown to inhibit *Escherichia coli*.^8–11^ Blue honeysuckle, a cold-climate *Caprifoliaceae* shrub, produces anthocyanin- and flavonoid-rich berries with reported antibacterial activity against foodborne and human pathogens.^12,13^ Golden root, a circumboreal *Crassulaceae* used across Tibetan, Chinese, and Korean medicine, contains phenolic and glycoside compounds with reported bacteriostatic and bactericidal activity against both Gram-positive and Gram-negative bacteria.^14,15^ Nevertheless, these three extracts also span a wide range of chemical complexity and pigmentation, properties that complicate their evaluation by conventional methods.

Conventional 96-well plate microdilution and agar-diffusion minimal inhibitory concentration (MIC) assays are the gold standards for conducting antimicrobial experiments. They are however laborious, time-consuming, and require large reagent volumes.^16–18^ More importantly, they measure average behavior across large bacterial populations, masking phenotypic heterogeneity and rare tolerant or resistant subpopulations.^19,20^ This population-averaging effect is particularly limiting when evaluating complex natural matrices, such as plant-derived bioactive compounds, that can modulate growth in subtle and heterogeneous ways.^21,22^ Natural extracts pose a further challenge for these bulk assays: their intrinsic pigmentation and chemical complexity, can directly interfere with the optical density measurements on which conventional MIC assays rely.

Droplet-based assays address these limitations by partitioning individual cells into isolated nano-to picolitre water-in-oil microreactors that enable subpopulation-level detection and amplify local concentrations of secreted molecules.^16–19^,^23–27^ Combined with high-resolution imaging, they are suited for capturing early aggregation and biofilm-like dynamics.^28^ However, microfluidics demands specialized expertise and infrastructure, limiting its adoption in standard laboratories.^29^ Polydisperse droplets generated by simple vortex- or shaking-based emulsification offer a democratized alternative that retains compartmentalization, high-throughput parallelization, and resolution of population heterogeneity. ^28–31^

The potential of droplet-based assays nevertheless depends on efficient and accessible image analysis. EasyFlow, our previously validated droplet image analysis platform^31^ still posed challenges for novice users. In this workflow, we present EasyFlow 2.0 (EF2), which offers stable, containerized deployment, flexible data formats, expanded plotting, automated statistical thresholding for classification and MIC estimation, joint analysis of multiple files or experiments, detailed phenotypic classification, and LLM-assisted data exploration. Using plant extracts and *Escherichia coli*, we benchmarked the polydisperse droplet-based assay against standard 96-well plate microdilution (**Figure 1**). To our knowledge, this is the first workflow combining droplet-based assays with a single analysis platform for both MIC estimation and aggregation-phenotype profiling in antibacterial screening, with potential application to complex sample matrices beyond plant extracts.

**Figure 1.**
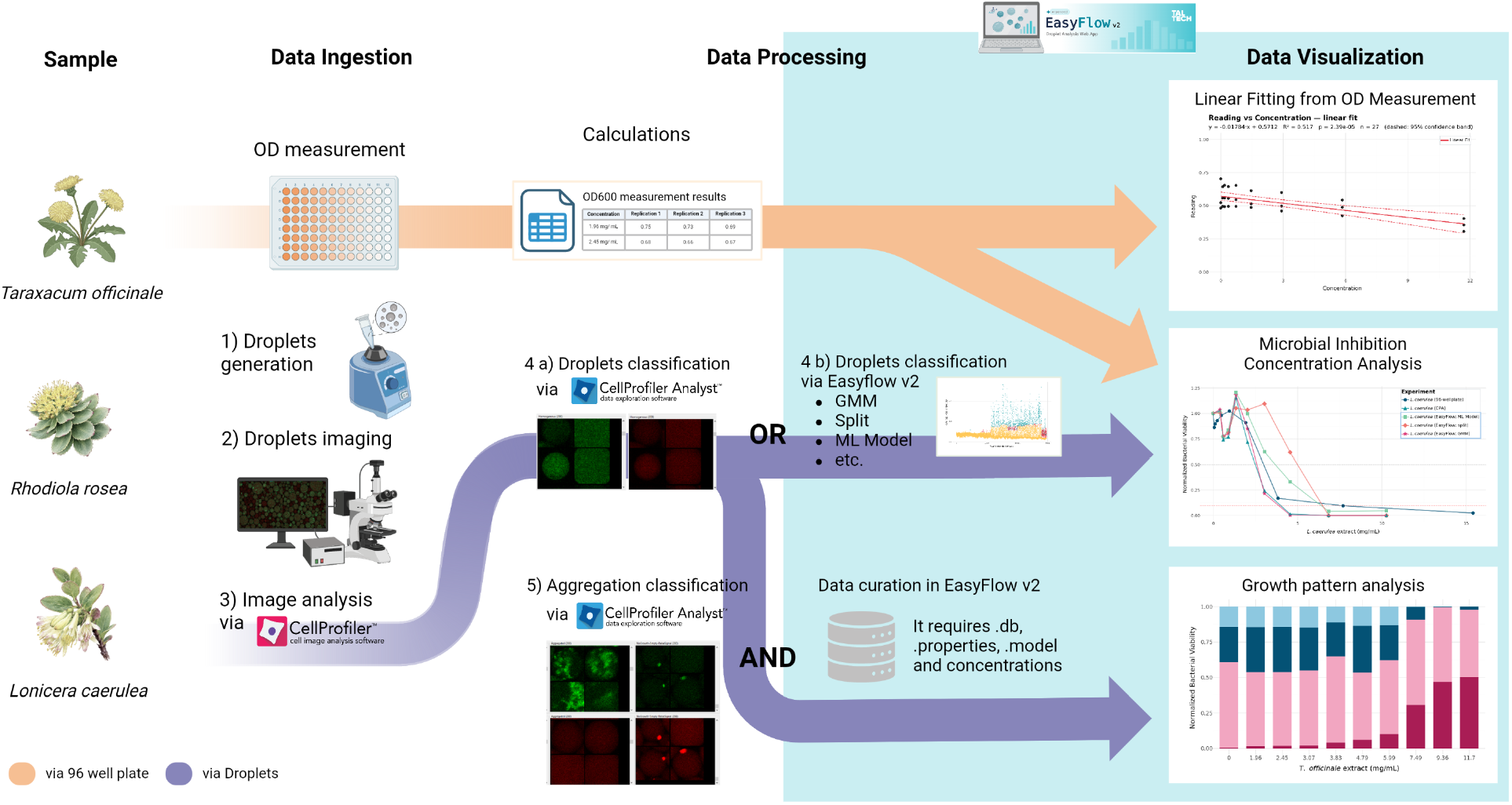
Study design benchmarking a polydisperse droplet-based assay against standard 96-well plate microdilution for minimal inhibitory concentration determination. Extracts of *Taraxacum officinale, Rhodiola rosea*, and *Lonicera caerulea* were chemically characterized by HPLC-MS and tested against *Escherichia coli* in parallel. In the 96-well plate microdilution assay, serial dilutions were plated and growth was quantified by optical density (OD^600^). In the droplet-based assay, extract, bacteria, and fluorescent dye were emulsified by vortexing, imaged by confocal microscopy, and segmented in CellProfiler™. *R. rosea* was excluded from droplet-based assay analysis because of extract-induced gelation. All other data were processed, visualized, and compared in EasyFlow 2.0 (EF2): linear fitting of OD^600^ against extract concentration (top), normalized viability versus extract concentration for 96-well plate microdilution and droplet-based assays (middle), and the proportion of droplets divided into four classification classes: “Homogenous,” “NoGrowth-Empty,” “Aggregated,” and “NoGrowth-Empty-FalseSignal” across extract concentrations (bottom).

## EXPERIMENTAL SECTION

### Plant extracts and chemical characterization

*Taraxacum officinale* flowers and *Rhodiola rosea* roots were collected from private gardens in Võrumaa and Saaremaa, Estonia, respectively, and dried *Lonicera caerulea* berries were purchased from Kingu Farm (Valgjärve, Estonia). Air-dried, ground material was extracted with 60% ethanol (v/v) at a 1:20 (w/v) ratio using shaking and ultrasound-assisted extraction. After centrifugation, ethanol was removed by vacuum evaporation, and the residue was redissolved in ultrapure water to give ethanol-free extracts at two-fold concentration. Soluble solid content was determined gravimetrically. Total polyphenols and total flavonoids were quantified in triplicate by Folin-Ciocalteu and AlCl_3_ colorimetric assays^32^, respectively using UV-Vis spectrophotometry against gallic acid and quercetin calibration standards and expressed as mg gallic acid equivalents (GAE) or quercetin equivalents (QE) per gram of plant material. Extracts were further analyzed by HPLC-DAD-MS/MS using a Poroshell 120 EC-C18 column with a water/acetonitrile gradient containing 0.1% formic acid, DAD detection at 254 nm, and negative-ion electrospray MS/MS over m/z 100-1000. Compounds were identified by comparison of MS/MS fragmentation patterns with literature. Full reagent sources, instrument specifications, calibration details, and chromatographic gradient parameters are provided in **Supplementary Methods**.

### 96-well plate microdilution assay and analysis

Conventional MIC assay was adapted from CLSI broth microdilution guidelines. Extract stock solutions (starting from dry matter concentrations found in **Supplementary Table S1**) were two-fold serially diluted across a 96-well plate in three technical replicates per extract, inoculated with exponential-phase GFP-labelled *Escherichia coli* JEK 1036^33^ at 5 × 10^5^ CFU/mL or LB broth-only controls, giving final concentrations of 23.4 to 0.046 mg/mL for *T. officinale*, 30.8 to 0.060 mg/mL for *L. caerulea*, and 52.2 to 0.102 mg/mL for *R. rosea*. Plates were incubated overnight at 37°C. The assay was performed in three independent experiments. Optical density at 600 nm (OD^600^) was measured using a Tecan GENios Pro plate reader (Tecan, Austria GmbH). Visual overview of the method is provided in **Supplementary Figure S1**. Background absorbance was corrected by subtracting values from extract-containing wells without bacteria from corresponding wells with bacteria and extract, following da Silva et al. ^34^ Corrected OD^600^ values were averaged across technical replicates within each independent experiment, and the resulting means (**Supplementary Excel S1)** were uploaded to EF2 and fitted by linear regression against extract concentration for each extract.

### Droplet-based assay and analysis

Polydisperse water-in-oil droplets were generated by vortexing equal volumes of aqueous phase (extract,

*E. coli* JEK 1036 at 5 × 10^5^ CFU/mL, and Alexa Fluor 647-dextran) and oil phase (Novec HFE 7500 fluorocarbon oil with PFPE-PEG-PFPE surfactant) for 5 seconds at maximum speed.^31,33^ Extracts were serially diluted in nine steps and mixed with an equal volume of bacterial suspension, giving final concentrations of 11.7 to 1.96 mg/mL for *T. officinale* and 10.27 to 0.4 mg/mL for *L. caerulea*, plus extract-free (water) control. Samples were incubated overnight at 37°C, then loaded into a Countess cell counting chamber slide (Invitrogen, Thermo Fisher Scientific) and imaged on a Zeiss LSM 900 confocal microscope (10×, 488/640 nm). Twenty images (mean 68 droplets/image; 0.56 nL mean droplet volume) were acquired per condition. *R. rosea* droplets were excluded from analysis due to extract-induced gelation and fluorescence loss (**Supplementary Figure S2A**). Droplets were segmented in CellProfiler™ 4.2.8 (CP) using Alexa Fluor 647 channel, adaptive Otsu thresholding, and shape filters. Size, shape, pixel intensity, and Haralick texture features were exported to either CellProfiler Analyst™ (CPA) or directly to EF2 (see EF2 analysis platform below). In CPA, 800 manually annotated droplets (200 per class, 100 from each extract) were used to train a Random Forest classifier model with four classes: “Homogenous,” “NoGrowth-Empty,” “Aggregated,” and “NoGrowth-Empty-FalseSignal”

(**Supplementary Figure S2B**). Classifier performance was evaluated by cross-validation on the annotated droplets in CPA, and the overall accuracy and per-class precision, recall, and F1-scores were calculated from the cross-validated predictions. The trained model from CPA was then further applied in EF2 for analysis of droplets. In total, 27,059 droplets from 400 images were analyzed. For the full analysis of the pipeline scheme, see **Supplementary Figure S3**.

### EasyFlow 2.0 (EF2) analysis platform

EF2, the updated version of EasyFlow ^31^, is deployed as a single-command Docker container (tested on macOS and Linux). It accepts Excel or CSV files as well as CP databases, properties files, and CPA models, and can host multiple files or experiments for joint analysis and visualization.

EF2 classifies droplets as positive or negative from intensity and texture features using five thresholding strategies: (1) a two-component Gaussian mixture model (GMM) fitted to the intensity distribution by expectation-maximization^35^ with the threshold placed at the posterior decision boundary (P(positive | intensity) = 0.5), an approach also used in digital PCR.^36–38^ and accepted only when Ashman’s D ≥ 2;^39^ (2) Otsu’s between-class variance;^40^ (3) a robust background cutoff (median + 3·MAD or Median Absolute Deviation)^41^ derived from the highest-concentration control; (4) a per-concentration split that thresholds each group independently, correcting baseline intensity drift between imaging sessions; and (5) a user-defined manual threshold set with a slider. When the GMM criterion is not met, the threshold reverts to Otsu’s method or to the robust background cutoff. EF2 can also train an experimental machine-learning classifier on the labels produced by the active strategy, using user-selected features. Unlike CPA, where the training set must be curated by manually sorting example objects, EF2 generates the training labels automatically: a stratified random sample that preserves the positive/negative ratio (capped at 50,000 droplets) is split 70/30 into training and test sets using a fixed seed for reproducibility, and the held-out portion estimates the agreement between the model and the threshold classification.

EF2 supports two routes for classifying droplets from CP output (**Supplementary Figure S3**), which can be provided as a database or as a per-object table (.csv or .xlsx). In the first route, the user classifies droplets directly in EF2 as positive or negative using one of the thresholding strategies above, without any labelled training data. For the second route, the user can supply a CPA model and database file, which EF2 can use to assign class labels to every droplet, enabling the four-class growth-pattern analysis (“Homogeneous”, “Aggregated”, “NoGrowth-Empty”, and “NoGrowth-Empty-FalseSignal”). Positive-droplet fractions were normalized to untreated control, and the MIC was defined as the lowest concentration at which normalized viability fell to ≤10% (MIC50: ≤50%). The same thresholds were applied to the normalized OD^600^ data from the 96-well plate microdilution assay. OD^600^ data from the plate reader can be uploaded alongside droplet data and fitted by linear regression (**Figures 2)**. 96-well plate microdilution and droplet-based assay results can be compared side by side on a shared concentration axis (**Figures 3**).

**Figure 2.**
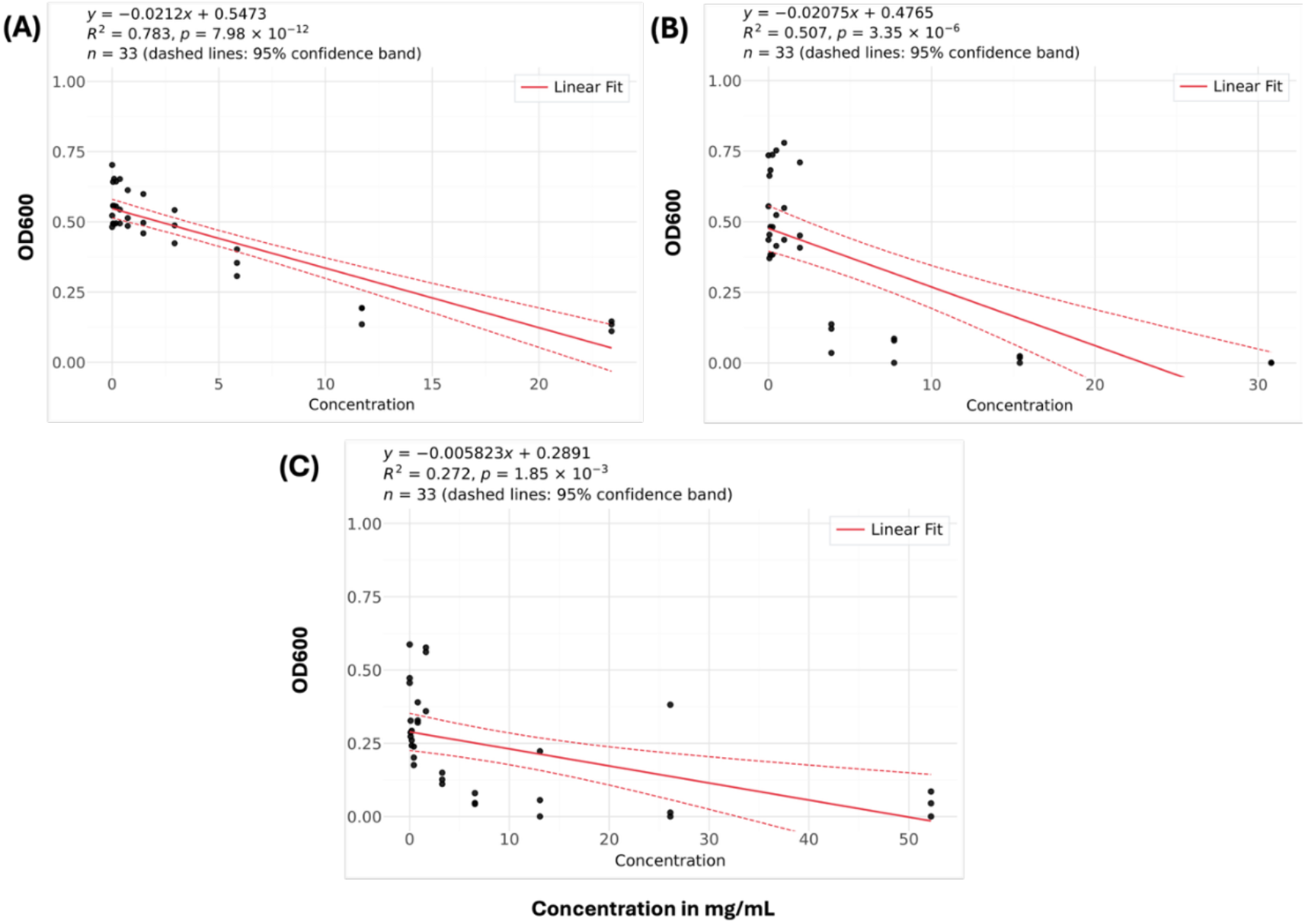
Dose-dependent decrease in *E. coli* growth in 96-well plate microdilution assay. Background-corrected OD^600^ is plotted against extract concentration (mg/mL) for (**A**) *Taraxacum officinale*, (**B**) *Lonicera caerulea*, and (**C**) *Rhodiola rosea*. Each point represents the mean of six technical replicates from one independent experiment (33 points per panel: 11 concentrations, including the extract-free control, across three independent experiments). Solid red lines show least-squares linear fits, and dotted lines indicate 95% confidence bands. Slopes were significantly negative for all three extracts (A: R^2^ = 0.783, p = 7.98 × 10−^12^; B: R^2^ = 0.507, p = 3.35 × 10−^6^; C: R^2^ = 0.272, p = 1.85 × 10−^3^). Fits were generated in EF2.

**Figure 3.**
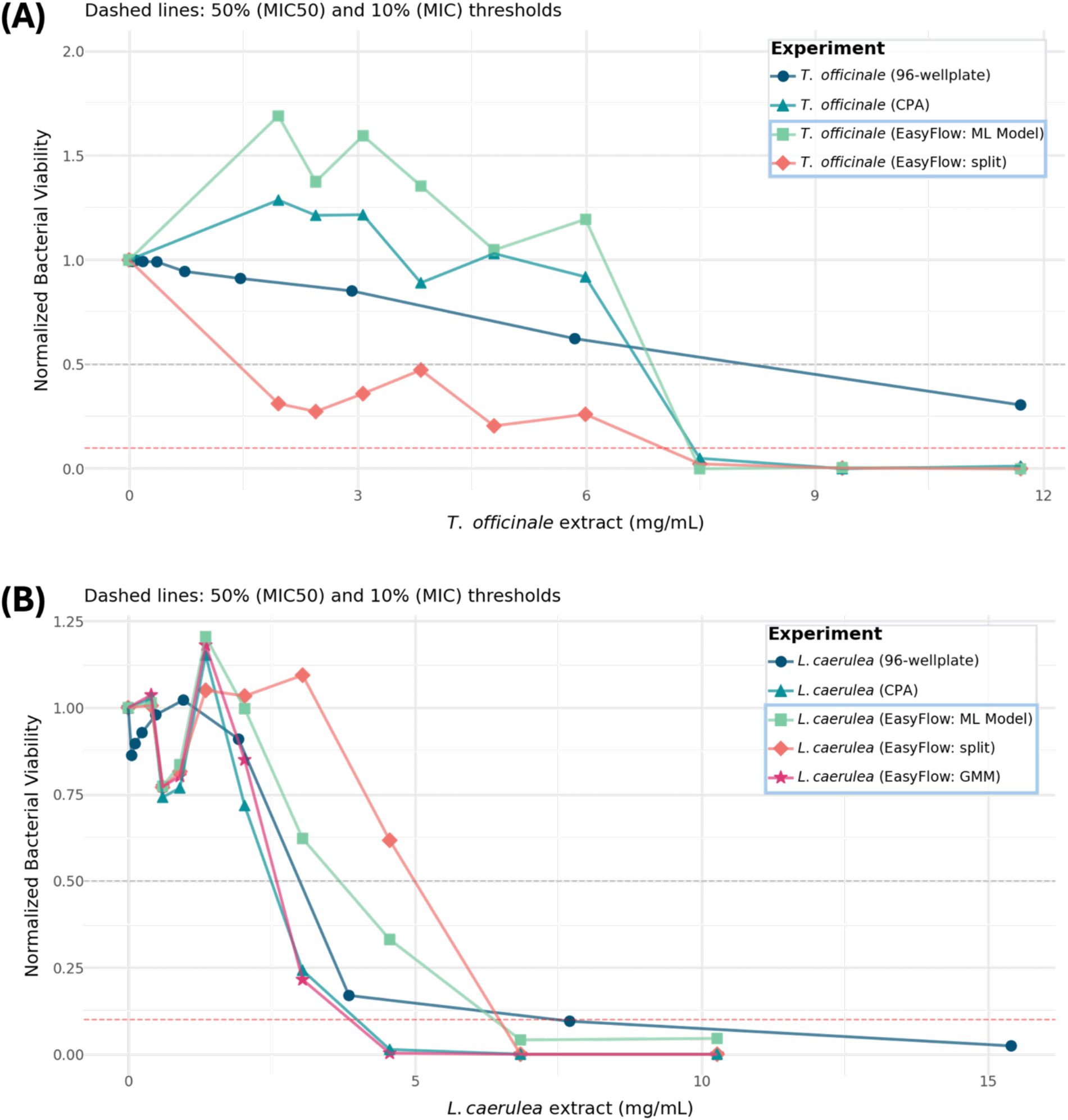
Droplet-based classification methods influence viability estimates and MIC determination. Normalized *E. coli* viability after treatment with (**A**) *Taraxacum officinale* and (**B**) *Lonicera caerulea*, measured by 96-well plate microdilution (background-corrected OD^600^) and droplet-based assay (fraction of positive droplets). CellProfiler™ outputs were classified using EF2 per-concentration split thresholding, a machine-learning (ML) classifier trained on threshold-derived labels, or a gated two-component Gaussian mixture model (GMM) and compared with a CellProfiler Analyst (CPA) Random Forest model. GMM results are shown only for *L. caerulea*, as the droplet intensity distributions for *T. officinale* lacked clear population separation. Values are normalized to untreated control (1.0), and concentrations are final assay concentrations. Microdilution results are shown up to 11.7 mg/mL for *T. officinale* and 15.4 mg/mL for *L. caerulea*; complete series are provided in Methods. Dashed lines indicate the 50% (MIC^50^) and 10% (MIC) viability thresholds. ML-derived estimates broadly followed CPA results, whereas threshold-based estimates varied, particularly for *T. officinale*, consistent with overlapping growth and background signals. Droplet-based assay reached MIC at lower extract concentrations than microdilution for both extracts.

EF2 also includes an LLM-assisted assistant for querying data and generating basic visualizations and transformations. Requests are evaluated through a whitelisted interpreter rather than executed as code, which removes the code-injection risk,^42^ and only summary statistics are sent to the language model which can be a local open-weight model (Ollama^43^) or other third-party providers. EF2 runs on FastAPI with plotnine for plotting.^44^ Additional modules not used in this study are described in **Supplementary Figure S3**. The EF2 repository is currently available upon request, and a password for direct testing at https://easyflow.taltech.ee is provided to interested researchers.

## RESULTS AND DISCUSSION

### Plant extracts showed distinct phytochemical profiles and differed in phenolic content

Colorimetric and HPLC-DAD-MS/MS analyses showed that *R. rosea* root extract contained the highest, and

*T. officinale* flower extract the lowest, concentrations of total polyphenols and flavonoids, with a strong positive correlation between phenolic content and soluble solid content across all three extracts

(**Supplementary Table S1, Supplementary Figure S4**). Compound identification (**Supplementary Figures S5-S7**) confirmed extract-specific phytochemical profiles consistent with previous reports: *T. officinale* was dominated by hydroxycinnamic acid derivatives (including chicoric acid) and luteolin glycosides, ^45,46^

*L. caerulea* by kaempferol- and taxifolin-hexoside isomers, identified from the characteristic m/z 285 kaempferol aglycon fragment, together with the iridoid glycoside loganic acid ^47,48^ loganin 7-O-pentoside, and rutin (anthocyanins, though commonly reported in this species, were not detected under these conditions), and *R. rosea* by prodelphinidin derivatives, sachaloside, rhodioloside, and rosarin/rosavin isomers. ^14,49^ This chemical diversity, spanning phenolic content, pigmentation, and soluble solids, provided astringent, chemically heterogeneous test case for the proposed droplet-based workflow.

### Pigmentation complicated analysis of *E. coli* growth inhibition in 96-well plate microdilution assays

In the 96-well plate microdilution assay, increasing concentrations of *T. officinale* and *L. caerulea* reduced *E. coli* growth in a dose-dependent manner (**Figure 2A, B**). *T. officinale* reduced OD^600^ but did not reach the MIC within the tested range, whereas *L. caerulea* reached the MIC at 7.70 mg/mL **(Supplementary Excel S1**). For *R. rosea*, the OD^600^ response was irregular and differed between independent experiments, with OD^600^ reduced even at some of the lowest concentrations (**Figure 2C**). The extracts differed in intrinsic pigmentation, which was strongest for *R. rosea* and *L. caerulea*, consistent with their higher phenolic and soluble-solid content (**Supplementary Figure S8**). This color likely interfered with OD^600^ readings even after background correction, limiting the reliability of this bulk readout.

### Droplet-based assays reached MICs at lower extract concentrations than 96-well plate microdilution

At high extract concentrations, two extract-specific imaging artifacts were observed that affected droplet-based image analysis. *R. rosea* extracts formed irregular, semi-solid droplet aggregates with uneven or blocked fluorescence, consistent with the extract’s high starch content, prodelphinidins, and glycosides (rosavin, rosarin, salidroside), which promote hydrogen bonding and increased viscosity upon solvent removal (**Supplementary Figure S2A, S7**).^14,50 51,52^ *T. officinale* and *L. caerulea* extracts produced bright fluorescent clumps visible in both Alexa Fluor™ 647 and GFP channels, reflecting extract-induced aggregation of the added fluorescent dye (**Supplementary Figure S2**).^51,53^

Fluorescence images of the droplets were segmented in CP, and the per-object feature tables were exported as .xlsx files **(Supplementary Excel S2 and S3)** and loaded into EF2. Within EF2, the data were grouped by concentration metadata and explored with LLM assistance, for example to normalize the average pixel intensity via prompt (**Supplementary Figure S11**). Droplets were classified as positive (growth) or negative (empty or no growth) from intensity and texture features using the thresholding strategies described in Methods; the manual threshold was not used because it is subjective and can differ between users. A machine-learning classifier trained on the labels from the active thresholding strategy was also applied. Positive-droplet fractions were then used to estimate the MIC in the EF2 MIC module (**Figure 3)**. For both *T. officinale* and *L. caerulea*, the proportion of positive droplets decreased with increasing extract concentration, and the 96-well plate microdilution and droplet-based assay results were compared (**Figure 3**).

### Droplet-based assays revealed extract-specific effects on bacterial aggregation

EF2’s automated thresholding methods classify droplets only as positive or negative. This is sufficient for viability and MIC estimation but cannot resolve growth phenotypes or separate bacterial growth from fluorescent artifacts. A Random Forest classifier was thus trained in CPA to sort droplets into four classes: “Homogeneous” (growth with an evenly distributed signal), “Aggregated” (growth with a clustered signal), “NoGrowth-Empty” (no growth), and “NoGrowth-Empty-FalseSignal.” The last class captures empty droplets containing the dye clumps described above. Because bacterial growth is visible only in the GFP channel, whereas these artifacts appear as small, equally intense regions in both fluorescence channels, the two can be told apart, and assigning the artifacts to a dedicated class prevented them from inflating either viability or aggregation estimates (**Supplementary Figure S2**). The classifier achieved an overall cross-validated accuracy of 88.6% (**Supplementary Figures S9**), with per-class F1-scores of 0.85-0.93 (**Supplementary Figures S10**). The per-droplet classifications were exported as CSV and loaded into EF2, which generated the viability and growth-pattern statistics and visualizations shown in **Figures 3 and 4**.

**Figure 4.**
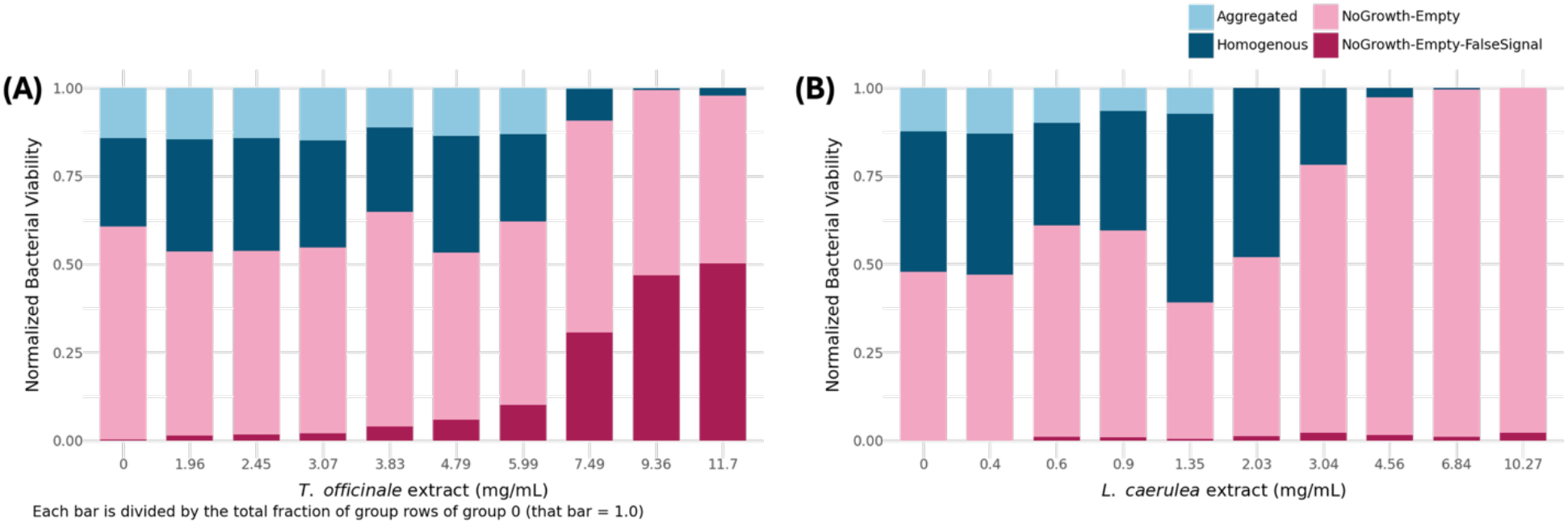
Effect of plant extracts on *E. coli* growth patterns in the droplet-based assay. Stacked bars show the fraction of droplets classified in EF2 using the CPA Random Forest model at each final extract concentration: bar segments are shaded in light and dark blue for Aggregated and Homogenous, while light and dark raspberry pink for NoGrowth-Empty and NoGrowth-Empty-FalseSignal, respectively. (**A**) *Taraxacum officinale*: the aggregated fraction remained approximately constant up to 5.99 mg/mL. At 7.49 mg/mL and above, where the MIC was reached, almost all droplets were classified as no growth or false signal. (**B**) *Lonicera caerulea*: the aggregated fraction decreased with increasing extract concentration and was absent at 2.03 mg/mL and above, whereas homogeneous growth persisted up to 3.04 mg/mL.

Both *T. officinale* and *L. caerulea* extracts inhibited *E. coli* in a concentration-dependent manner in the 96-well plate microdilution and droplet-based assays (**Figure 3**). In the droplet-based assay, the classification method altered the assignment of individual droplets as positive or negative, yet all methods yielded the same overall trend of decreasing viability with increasing extract concentration. Because droplet classification inherently involves a degree of subjectivity, the CPA model trained on manually annotated droplets was used as the reference method. Based on this classification, the droplet-based assay reached the MIC at lower extract concentrations than 96-well plate microdilution for both extracts: 7.49 mg/mL for *T. officinale*, for which the 96-well assay did not reach the MIC within the tested range, and 4.56 mg/mL for *L. caerulea*, compared with 7.70 mg/mL in the 96-well assay. At lower extract concentrations, the results of the EF2 threshold-based methods varied, particularly for *T. officinale* (**Figure 3A**). This is likely because overlapping signal distributions and dye clumps make the placement of a single intensity threshold less reliable, whereas the CPA model assigns false-signal droplets to a separate class. Normalized viability values above 1.0 at low concentrations likely reflect biological variability and classification uncertainty rather than growth stimulation.

Growth-pattern classification of droplets (**Figure 4A, B**) showed that increasing *L. caerulea* concentration reduced the fraction of aggregated droplets, until none were left from 2.03 mg/mL, while homogeneous growth was still present at 3.04 mg/mL (**Figure 4B**). In contrast, the fraction of aggregated droplets in *T. officinale* was stable up to 5.99 mg/mL and disappeared only at 7.49 mg/mL and above, where growth was inhibited and most droplets were classified as “NoGrowth-Empty” or “NoGrowth-Empty-FalseSignal” (**Figure 4A**). *T. officinale* therefore reduced viability without changing bacterial aggregation. Finally, the most informative features for the CPA Random Forest classifier were Haralick texture features, particularly Sum Variance. This agrees with our previous droplet-based analysis^28^, and highlights that polydisperse droplet-based assays combined with CP feature measurements can facilitate determination of growth phenotypes.

The strong pigmentation of *L. caerulea* (**Supplementary Figure S8**) likely caused optical interference in the 96-well plate microdilution assay even after background subtraction, a limitation that the image-based droplet assay reduces by classifying fluorescence patterns rather than bulk absorbance. The lower MIC values in droplets are also consistent with reports that droplet-based assays often detect lower MIC values than bulk assays, which has been attributed to compartmentalization of single cells or small subpopulations reducing population-level protective mechanisms such as quorum sensing and collective stress responses. ^54,55^ Differences in the number of cells per droplet compared with per well may also contribute. Collectively, these results show that combining CPA droplet classification with EF2’s visualization and statistics resolves extract-specific artifacts and yields MIC and growth-pattern information from the same dataset. Supervised four-class classification in EF2 still requires a model trained in CPA, but EF2 supports reproducible analysis from the CP database, model, and output files alone, and its unsupervised thresholding provides automated viability and MIC estimates when no labelled training data are available.

## CONCLUSION

This study demonstrates a polydisperse droplet-based workflow centered on EasyFlow 2.0 (EF2) for evaluating the antibacterial activity and growth-pattern modulation of plant extracts without requiring microfluidic equipment. Using the CPA Random Forest model as the reference, the droplet-based assay reached the MIC at lower extract concentrations than 96-well plate microdilution. Classification of 27,059 droplets revealed extract-specific growth phenotypes: *L. caerulea* reduced *E. coli* aggregation, whereas *T. officinale* reduced viability without altering aggregation. Although all classification methods showed concentration-dependent growth inhibition, differences in droplet assignment highlighted the limitations of threshold-based classification when signal distributions overlap or fluorescent artifacts are present.

Extract-induced gelation prevented analysis of *R. rosea*, while dye aggregation in the other extracts required a dedicated false-signal class, underscoring the need for extract-dependent controls. EF2 supports automated viability and MIC estimation through unsupervised thresholding and, with CP measurements and a CPA-trained model, four-class growth-pattern analysis. To our knowledge, this is the first workflow combining droplet-based assays with a single analysis platform for both MIC estimation and aggregation-phenotype profiling in antibacterial screening of plant extracts. Future work should refine extract preprocessing, standardize droplet classification, and evaluate additional microorganisms and complex sample matrices.

## Supporting information

https://easyflow.taltech.ee

## AUTHOR CONTRIBUTIONS

I.S., S.B., M.V. contributed to conception and design. M.V., P.P., S.B., P.S-R., K.R. performed experiments. I.S., M.V., S.B., P.P., A.A.S. performed data analysis. All authors contributed to writing and reviewing the manuscript. All authors have read and agreed to the published version of the manuscript.

