## Supplementary material for "A droplet-to-data workflow for antimicrobial screening of complex matrices": https://easyflow.taltech.ee

### Supplementary Methods

#### Botanical material and extraction procedure

*Taraxacum officinale* flowers and *Rhodiola rosea* roots were collected from private gardens in Võrumaa and Saaremaa, Estonia, respectively. *Lonicera caerulea* dried berries were purchased from Kingu Farm (Valgjärve, Estonia). All materials were air-dried and stored at ambient temperature in the dark prior to extraction. Plant material was ground using a Bomann KSW 445 CB coffee grinder (China) and extracted with 60% ethanol (v/v) at a 1:20 plant material-to-solvent ratio (w/v) by shaking for 30 min at 250 rpm, followed by sonication at 640 W and 350 kHz for 30 min at 40 °C in a Sonorex™ Digital 10P bath (Bandelin, Germany). Extracts were centrifuged at 8000 rpm, ethanol was removed using a Laborota 4000 vacuum evaporator (Heidolph, Germany), and solid residue was dissolved in ultrapure water to produce two-fold concentrated, ethanol-free extracts. Soluble solid content was determined gravimetrically by drying samples to constant weight in a vacuum concentrator (Eppendorf, Germany).

#### Chemicals

Ultrapure water (≥18 MΩcm) was produced in-house using a Milli-Q water purification system (Merck KGaA, Darmstadt, Germany) and used to prepare all aqueous solutions. Extraction solvent ethanol (96.7%) was obtained from Sigma-Aldrich (Munich, Germany). Acetonitrile (≥99.9%) for HPLC eluents was purchased from Honeywell (Seelze, Germany), and formic acid (≥99.0%) from Fisher Chemical (Czech Republic). Total polyphenolic content was evaluated using 2 M Folin-Ciocalteu reagent (Sigma-Aldrich, Switzerland) and sodium carbonate (Sigma-Aldrich, Germany). Standard solutions were prepared from gallic acid monohydrate (Sigma-Aldrich, China) and 96.4% ethanol (Sigma-Aldrich, Germany). For total flavonoid detection, aluminium chloride was obtained from Fluka (Switzerland), and standard solutions were prepared from quercetin (≥99.0%, Lachema/Chemapol) and methanol (≥99.9%, Honeywell, Charlotte, NC, USA).

#### Colorimetric assays

Total polyphenols and total flavonoids were quantified using the Folin-Ciocalteu and AlCl3 methods, respectively. All colorimetric analyses were performed on a Varian Cary 50 Bio UV-Vis spectrophotometer (Agilent Technologies, Santa Clara, CA, USA). Total polyphenolic content was determined from calibration solutions of 10, 25, 50, 75, and 100 mg/L gallic acid in ethanol, prepared from a 5 g/L stock solution. Total flavonoid content was determined from calibration solutions of 2, 5, 10, 20, and 40 mg/L quercetin in methanol, prepared from a 2 g/L stock solution. All samples were measured in triplicate, and results are expressed as mg gallic acid equivalents (GAE) or quercetin equivalents (QE) per gram of plant material (mg GAE/QE/g) ± standard deviation.

#### HPLC-MS analysis

Before HPLC-DAD-MS/MS analysis, extracts were centrifuged, and the supernatant was diluted 5- or 10-fold (v/v) with ultrapure water and spiked with bicalutamide as internal standard. The injection volume was 5 µL. Separations were performed on an Agilent 1260 Infinity II system (Agilent Technologies, Santa Clara, CA, USA) equipped with an Agilent Poroshell 120 EC-C18 column (2.7 µm, 4.6 × 100 mm) maintained at 28 °C. The mobile phase consisted of ultrapure water (A) and acetonitrile (B), both containing 0.1% (v/v) formic acid. The gradient elution program was as follows: 5–50% B (0–20 min), 50–95% B (20–25 min), isocratic 95% B (25–30 min), 95–5% B (30–30.01 min), and isocratic 5% B (30.01–35 min). The flow rate was 0.6 mL/min. Detection was carried out with a DAD at 254 nm, with spectra recorded from 200 to 400 nm. MS/MS analysis was performed on an LC/MSD Trap XCT mass spectrometer (Agilent Technologies) with an electrospray ionization source, operating in negative-ion mode over m/z 100–1000. Nitrogen was used as nebulizing and drying gas, and helium as collision gas. Compounds were identified from MS/MS fragmentation patterns (automatic settings) by comparison with relevant literature.

**Supplementary Figures and Tables**

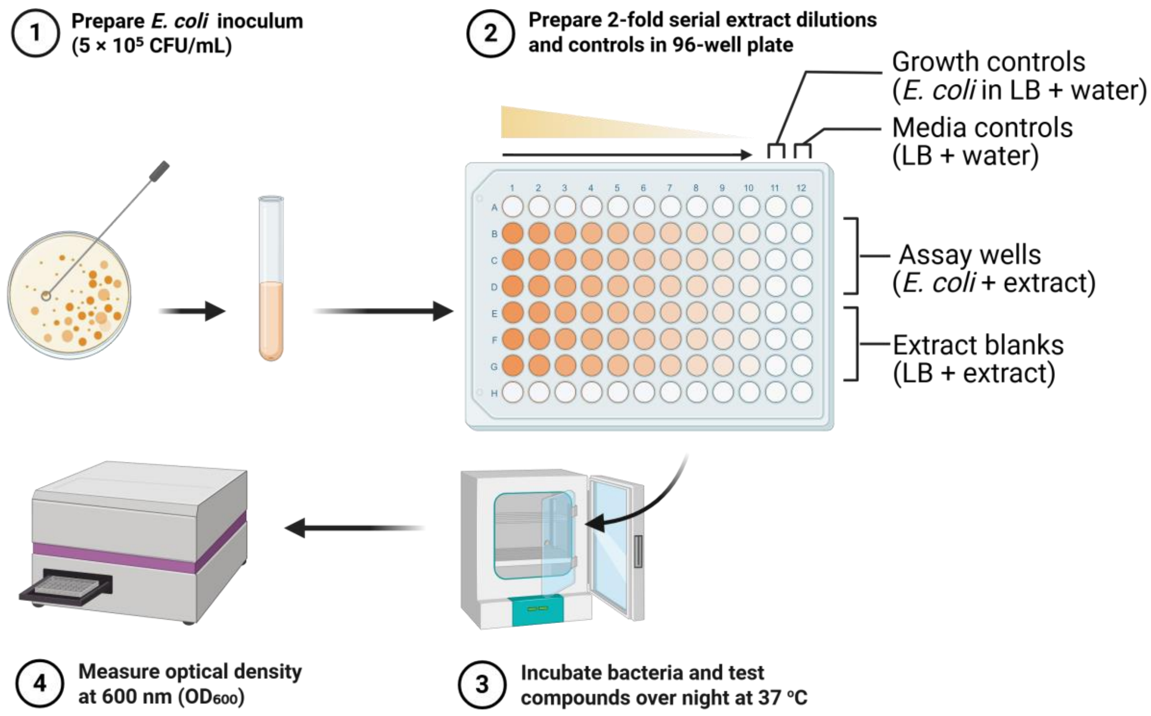


**Supplementary Figure S1**. Schematic layout and steps of the 96-well plate microdilution (MIC) assay used to evaluate the antibacterial activity of plant extracts. (1): *Escherichia coli JEK1036* culture at 5 × $10^5$ CFU/mL was prepared. (2): Rows B-G contained six technical replicates of each extract (*Taraxacum officinale*, *Rhodiola rosea*, and *Lonicera caerulea*), prepared as a two-fold serial dilution across the plate starting from stock solution in column 1. Following dilution, wells were inoculated with *E. coli* JEK1036 (5 × $10^5$ CFU/mL) or LB broth for negative controls. The final two columns contained no extract and served as growth and media controls. (3): Well plate was incubated over night at 37 °C. (4): Bacterial growth was quantified by measuring optical density at 600 nm ($OD_{600}$). To account for the intrinsic colour of the plant extracts (see **Supplementary Figure S8** for visual illustration), background absorbance was corrected using concentration-matched negative-control wells containing the corresponding extract dilution in LB broth without bacteria. This corrected $OD_{600}$ was used for MIC determination.

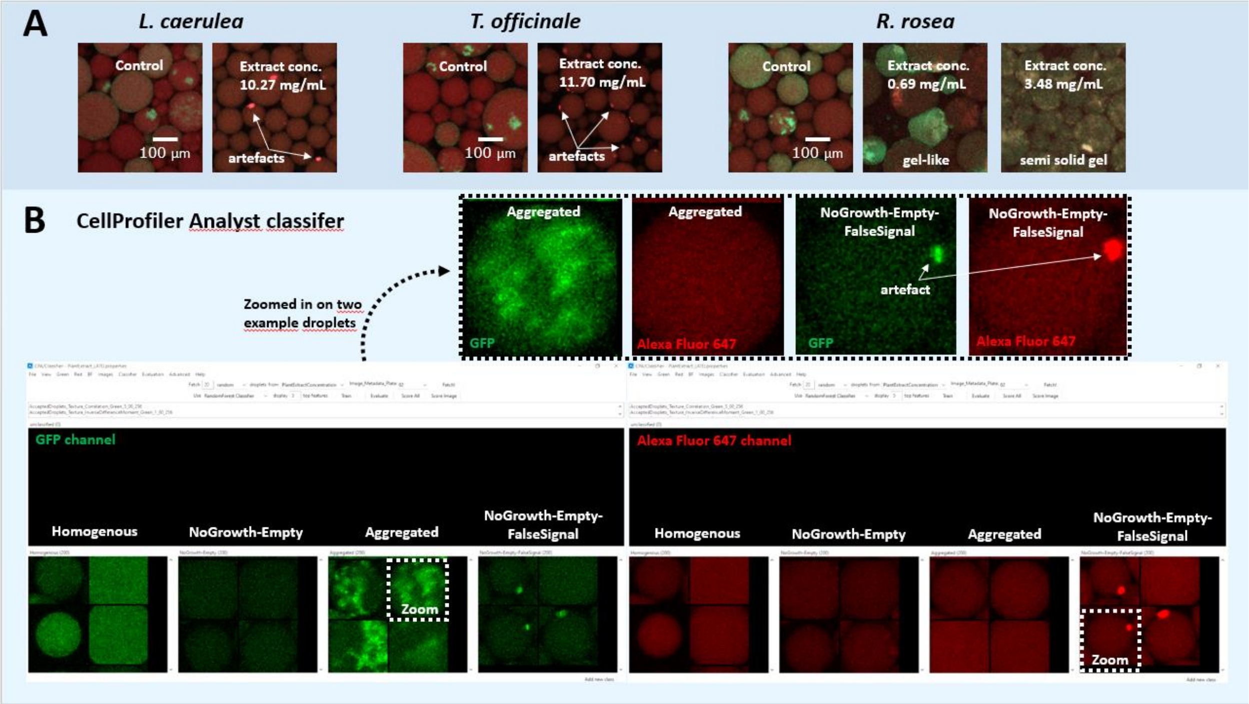


**Supplementary Figure S2**. **(A)** Example images of extract-induced fluorescence artifacts affecting droplet image analysis. High concentrations of *Rhodiola rosea* extract interfered with fluorescence emission and formed gel-like or semi-solid droplets, preventing reliable imaging and analysis and resulting in the exclusion of *R. rosea* from droplet-based analysis. For *Taraxacum officinale* and *Lonicera caerulea* extracts, high concentrations produced bright fluorescent clumps in the Alexa Fluor 647 channel, attributed to aggregation of insoluble plant components with the Alexa Fluor 647 dye. Because of their extremely high intensity, these aggregates were also visible in the GFP channel, where they could be mistaken for bacterial growth. True bacterial growth is detectable only in the GFP channel (GFP-labelled E. coli), whereas artifacts appear bright in both channels at the same position. **(B)** Illustration of how these artifacts were addressed during image analysis in CellProfiler Analyst. Example droplets are shown in the GFP (green) and Alexa Fluor 647 (red) channels for each class, with dotted boxes marking the zoomed regions. The Random Forest classifier was trained with a dedicated "NoGrowth-Empty-FalseSignal" class to prevent droplets containing artifacts from being misclassified as droplets with bacterial growth. Four classes were thus used in training: "Homogenous," "NoGrowth-Empty," "Aggregated," and "NoGrowth-Empty-FalseSignal."

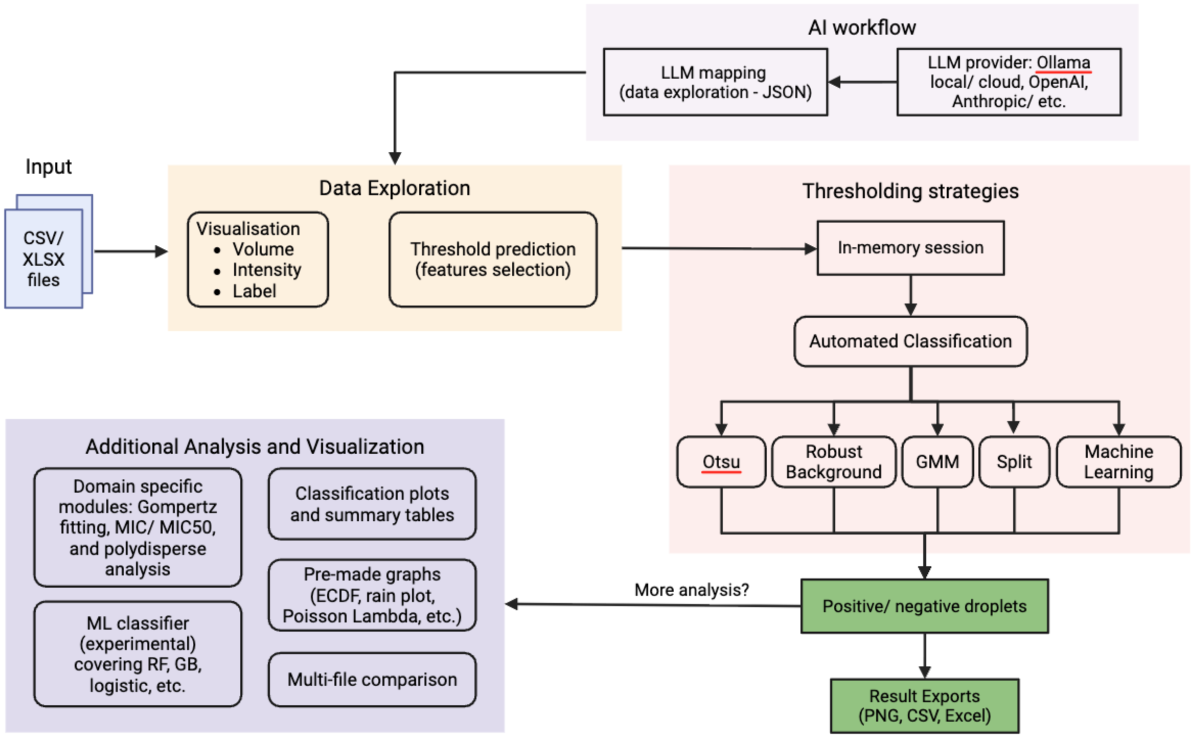


**Supplementary Figure S3.** Overview of EasyFlow v2 (EF2) platform design and user journey. Users upload one or more data files (CSV/XLSX) which contain output measurements from CellProfiler$^{TM}$ (CP) or 96-well plate microdilution optical density at 600 nm ($OD_{600}$) readings. CP measurements can then be used to perform visualization and data exploration while the $OD_{600}$ data can be used to compare result later in the analytical pipeline. In the data-exploration stage, the three columns that drive the analysis (droplet volume, signal intensity, and an optional grouping label) are recognized automatically or assigned by the user, and feature columns can be selected for the downstream machine-learning classifier. An optional AI workflow can also be enabled when uploading the data. If user use large-language-model provider (local or cloud Ollama, OpenAI, or Anthropic), the LLM will receive only aggregate summary statistics, leaving the raw droplet data never leave the server and returns machine-readable JSON (JavaScript Object Notation) data (column mappings, derived-column formulas, candidate feature sets). In the thresholding stage, droplets are classified as positive or negative per concentration group. The primary method is a two-component Gaussian Mixture Model fitted by expectation-maximization, with the threshold placed at the posterior decision boundary (P (positive | intensity) = 0.5), as established in digital PCR analysis and the fit is accepted only when the two populations are clearly separated (Ashman's D ≥ 2). When this criterion is not met, EF2 falls back to Otsu's method or to a robust background cutoff (median + 3·MAD) estimated from the highest antibiotic concentration group, and all thresholds remain user adjustable. The classified droplets feed the additional analysis and visualization modules, rendered with plotnine while classification plots and summary tables, pre-made views (ECDF, rain plot, Poisson concentration estimates), domain modules for Gompertz growth-curve fitting, MIC/MIC50 estimation, and polydisperse size-distribution analysis can be generated later in the platform. The side-by-side comparison of multiple files sharing experimental conditions, with non-parametric statistics (pairwise Mann–Whitney U tests, one-way ANOVA for more than two datasets) and an experimental machine-learning classifier (random forest, gradient boosting, or logistic regression) that evaluates whether user-selected feature columns are also available in the platform. User then can retrieve PNG figures and CSV/Excel tables for the result. EF2 is distributed as a Docker container for the bundled Ollama runtime) and runs fully offline when a local model is used.

**Supplementary Table S1.** Overview of the bioactive compound and soluble solid content concentrations of *T. officinale*, *R. rosea*, and *L. caerulea* extracts. Values are presented as mean ± SD (n = 3).

| Plant extract | Total polyphenols, mg GAE/mL | Total flavonoids, mg QE/mL | Dry matter, mg/mL |
|---|---|---|---|
| *T. officinale* | 0.99 ± 0.07 | 1.29 ± 0.70 | 46.80 ± 2.21 |
| *R. rosea* | 13.45 ± 1.43 | 19.99 ± 2.41 | 104.47 ± 5.60 |
| *L. caerulea* | 5.16 ± 0.40 | 5.46 ± 0.38 | 61.60 ± 3.12 |

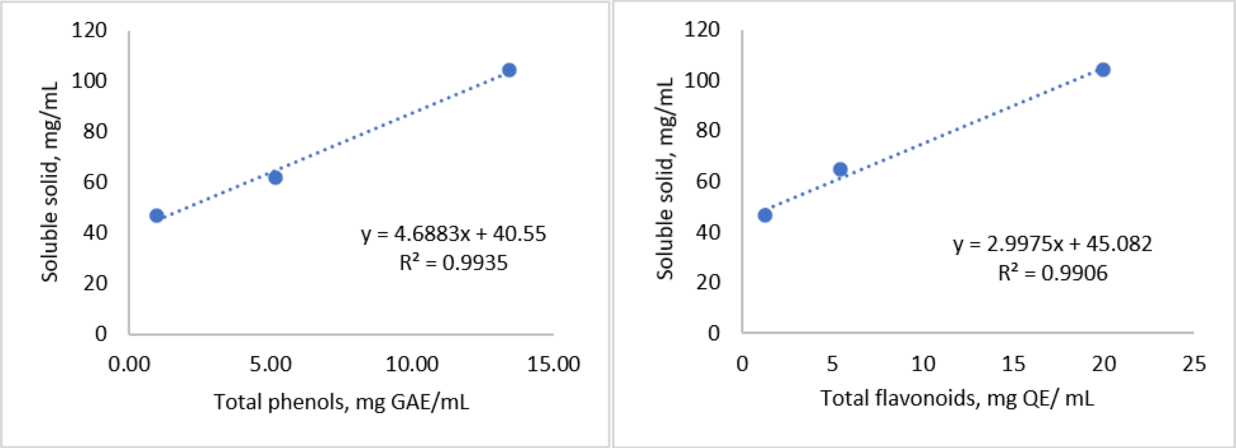


**Supplementary Figure S4**. Relationship between soluble solid content and total polyphenol (left) or total flavonoid (right) concentration in *Taraxacum officinale*, *Rhodiola rosea*, and *Lonicera caerulea* extracts. Linear regression equations and coefficients of determination ($R^2$) are shown.

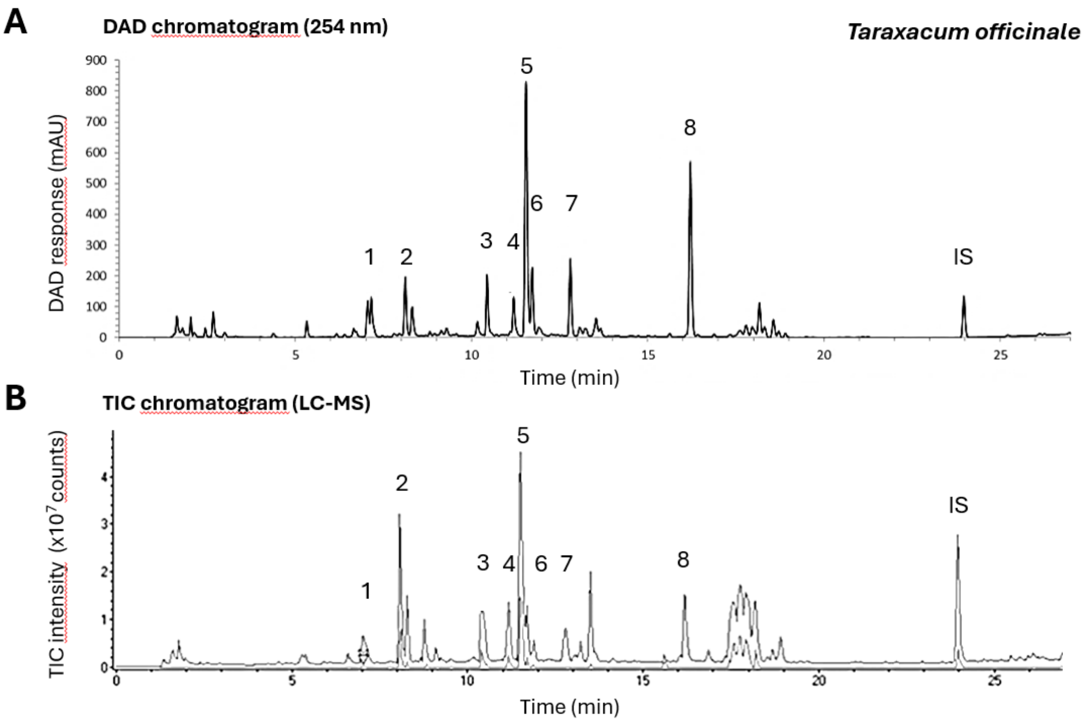


**Supplementary Figure S5**. Representative chromatograms of *Taraxacum officinale* flower extract obtained by (A) diode-array detection (DAD, 254 nm) and (B) liquid chromatography-mass spectrometry (LC-MS) total ion current (TIC). The numbered peaks correspond to: (1) *cis*-caftaric acid, (2) chlorogenic acid, (3) luteolin diglucoside, (4) luteolin-7-O-rutinoside, (5) chicoric acid, (6) luteolin-7-O-glucoside, (7) 3,5-di-O-caffeoylquinic acid, and (8) quercetin caffeoyl glucoside or a luteolin derivative. IS indicates the internal standard.

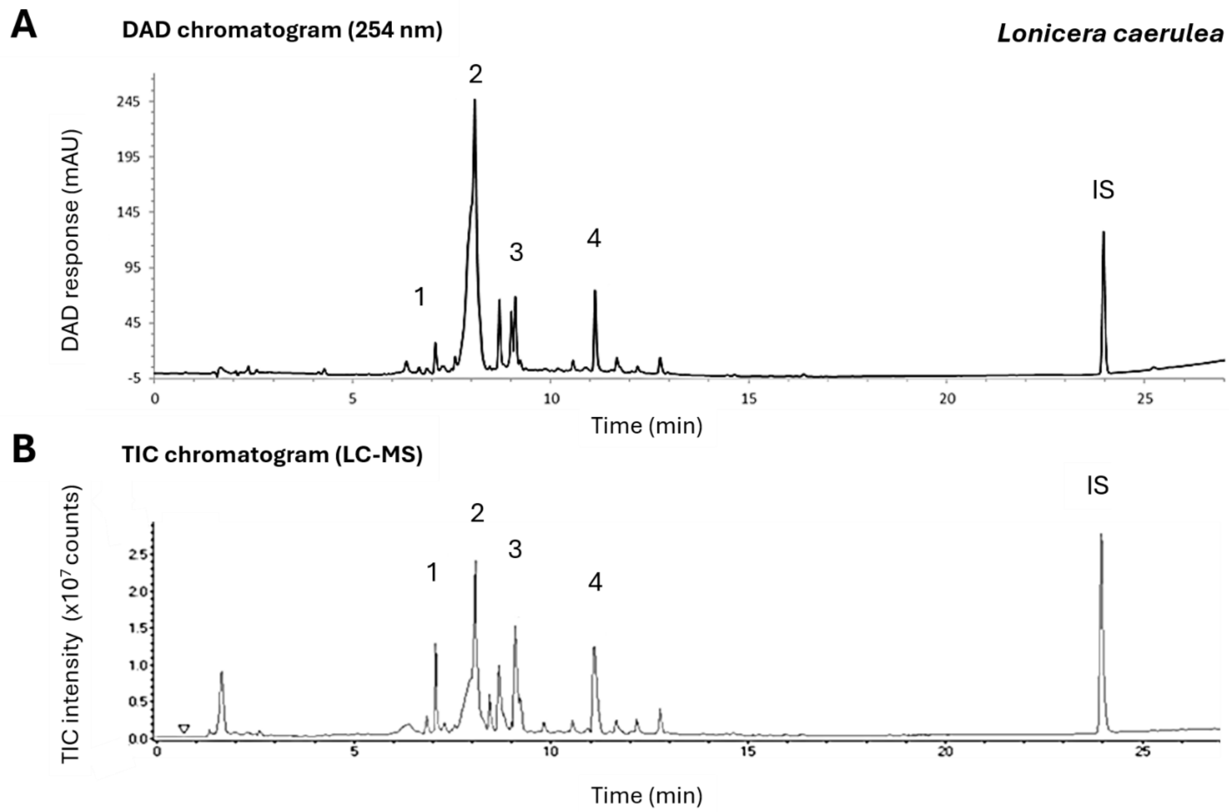


**Supplementary Figure S6**. Representative chromatograms of *Lonicera caerulea* L. berry extract obtained by (A) diode-array detection (DAD, 254 nm) and (B) liquid chromatography–mass spectrometry (LC–MS) total ion current (TIC). The numbered peaks correspond to: (1) loganic acid, (2) a mixture of kaempferol- and taxifolin-hexoside isomers, (3) loganin-7-O-pentoside, and (4) rutin. IS indicates the internal standard.

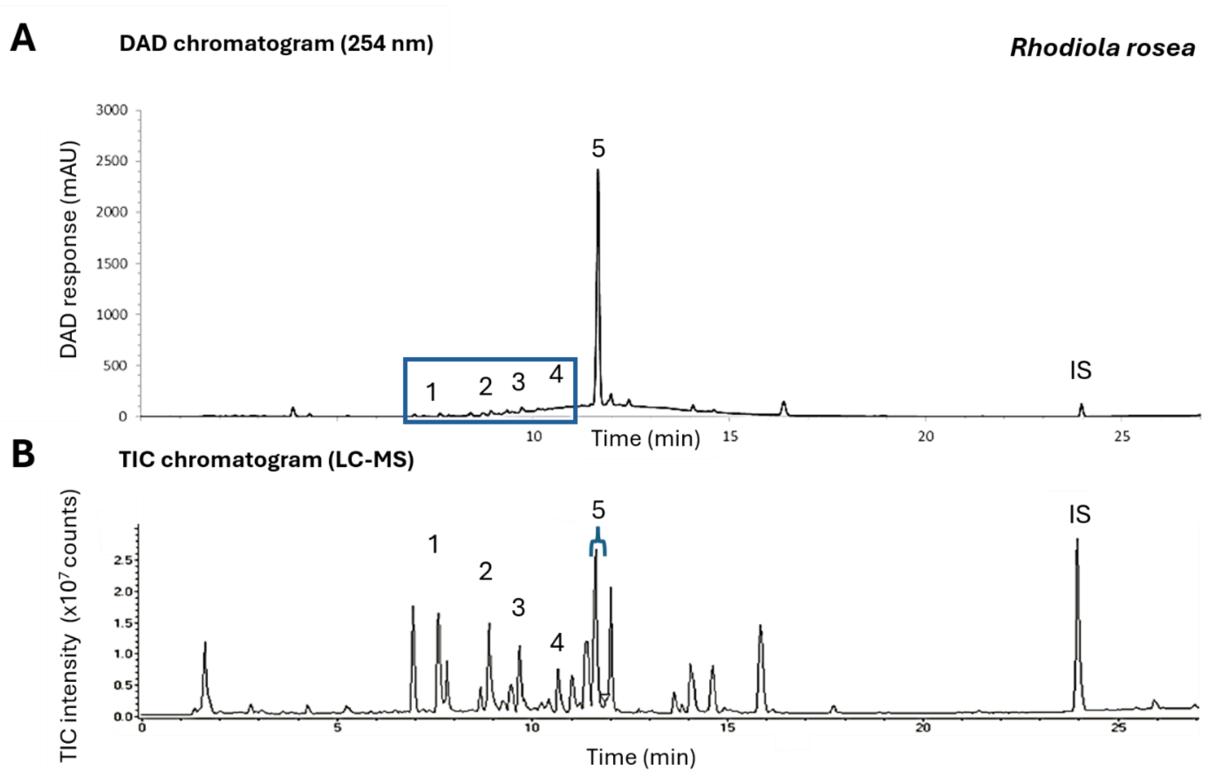


**Supplementary Figure S7**. Representative chromatograms of *Rhodiola rosea* root extract. (A) DAD chromatogram recorded at 254 nm. (B) Total ion chromatogram (TIC) obtained by LC–MS analysis. The numbered peaks correspond to: (1) prodelphinidin B-2-O-gallate, (2) prodelphinidin B-2-di-O-gallate, (3) sachaloside II/sacranoside A, (4) rhodioloside E, and (5) rosarin/rosavin. IS indicates the internal standard.

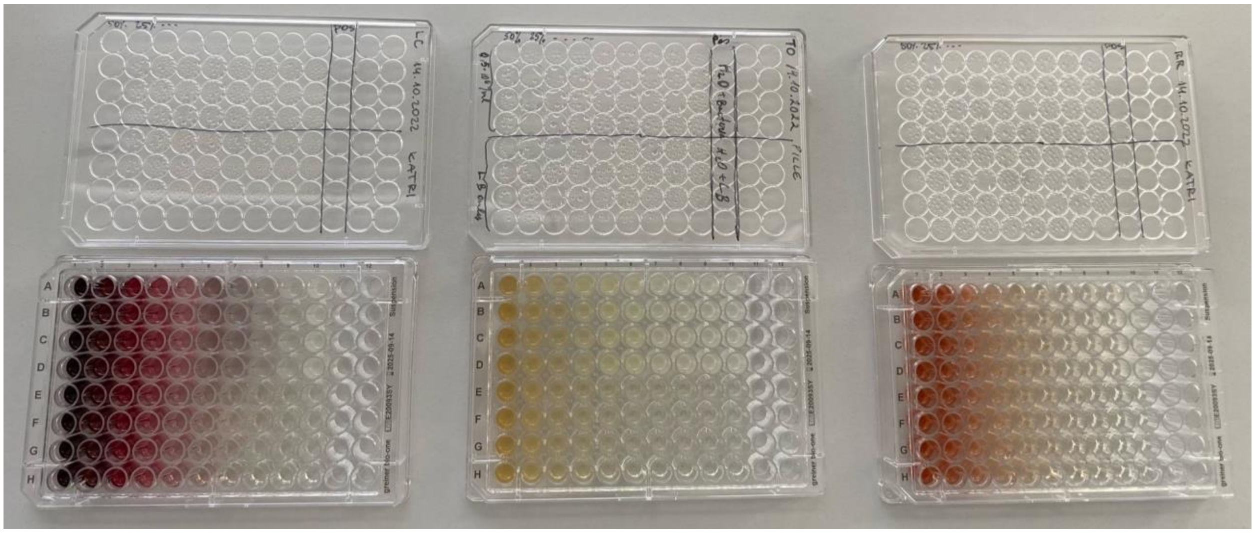

**Supplementary Figure S8.** Representative 96-well plate photos showing the intense colour of the respective plant extracts from left to right: *R. rosea*, *T. officinalis*, and *L. caerulea* respectively

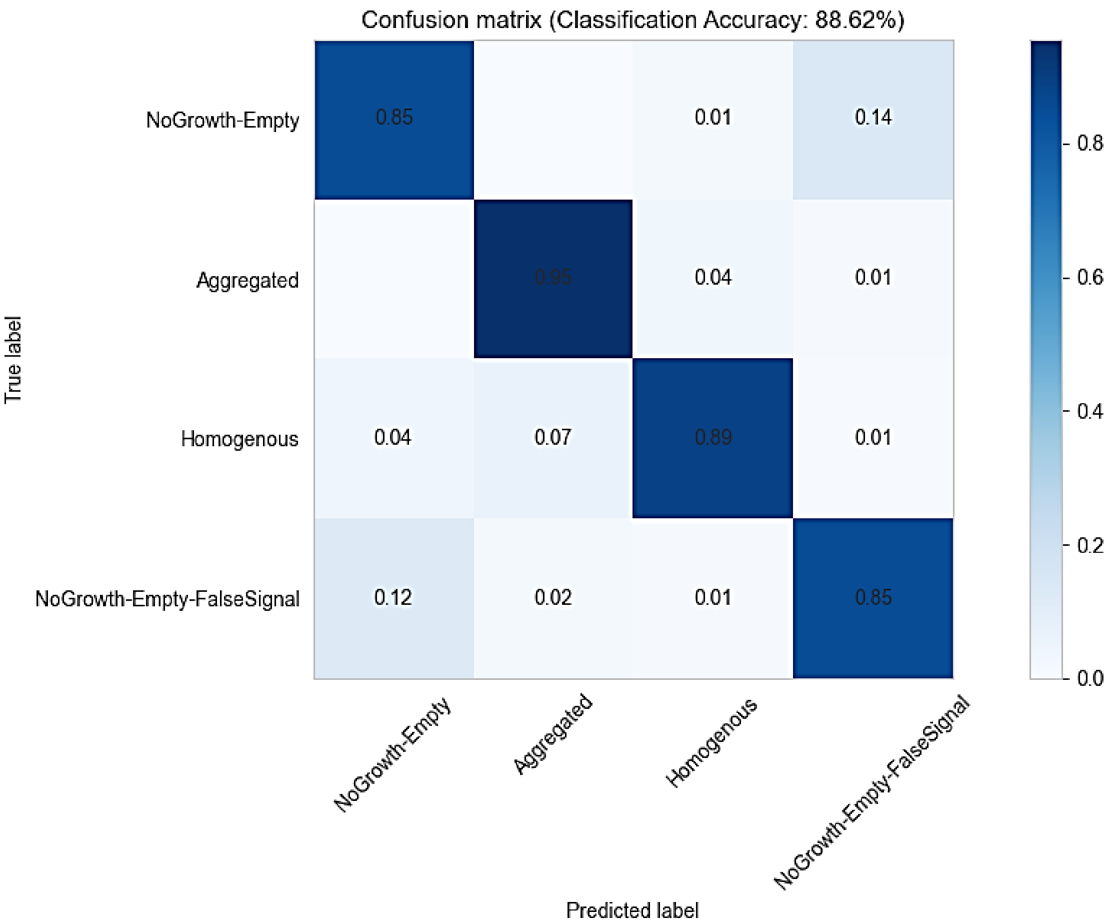


**Supplementary Figure S9**. Confusion matrix generated using the supervised machine-learning Classifier module in CellProfiler™ Analyst (CPA). Four droplet classes were manually defined within the Classifier: (i) "Homogenous", representing droplets with uniformly distributed bacterial growth; (ii) "NoGrowth-Empty" representing droplets with absent or minimal bacterial growth, including empty droplets; (iii) "Aggregated", representing droplets containing bacteria that formed localized, high-intensity aggregates; and (iv) "NoGrowth-Empty-FalseSignal", representing droplets with no bacterial growth but exhibiting highly fluorescent clumps caused by interaction between concentrated plant extract and Alexa Fluor™ 647, resulting in artifact-like "false" signals. The Classifier model was trained using 400 droplets per plant extract, randomly selected and manually annotated (100 droplets per class), yielding a combined training dataset of 800 droplets across both extract conditions. Classification was performed using the built-in Random Forest algorithm in CPA, leveraging the full set of features extracted from the CellProfiler™ pipeline. The confusion matrix summarizes the performance, classification accuracy of 88.62%, of the cross-validated model by comparing predicted class labels to the true manually assigned labels (training dataset) across the four predefined droplet classes.

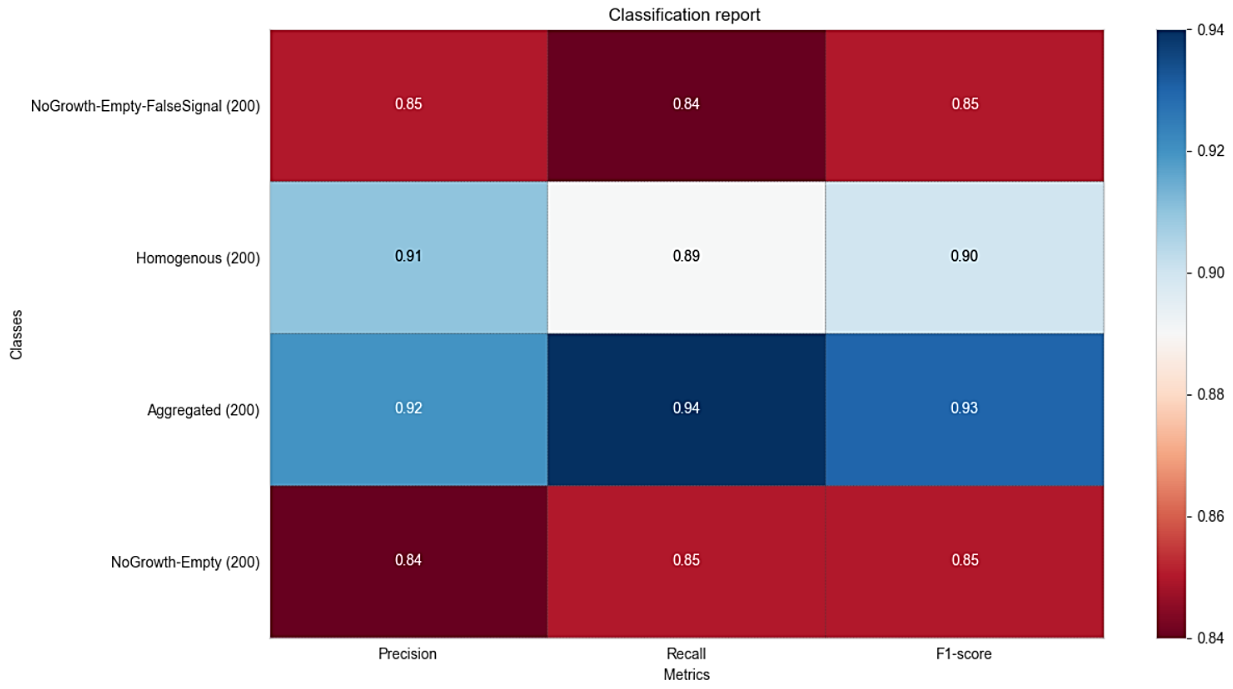


**Supplementary Figure S10**. Classification report generated using the supervised machine-learning Classifier module in CellProfiler Analyst™ (CPA). Four droplet classes were manually defined within the Classifier: (i) “Homogenous”, representing droplets with uniformly distributed bacterial growth; (ii) “NoGrowth-Empty” representing droplets with absent or minimal bacterial growth, including empty droplets; (iii) “Aggregated”, representing droplets containing bacteria that formed localized, high-intensity aggregates; and (iv) “NoGrowth-Empty-FalseSignal”, representing droplets with no bacterial growth but exhibiting highly fluorescent clumps caused by interaction between concentrated plant extract and Alexa Fluor™ 647, resulting in artifact-like “false” signals. The Classifier model was trained using 400 droplets per plant extract, randomly selected and manually annotated (100 droplets per class), yielding a combined training dataset of 800 droplets (total 200 droplets per class) across both extract conditions. Classification was performed using the built-in Random Forest algorithm in CPA, leveraging the full set of features extracted from the CellProfiler™ pipeline. The classification report summarizes the performance of the cross-validated model by comparing predicted class labels to the true manually assigned labels across the four predefined droplet classes. Performance metrics are reported as precision, recall metrics, and F1-score for each class.

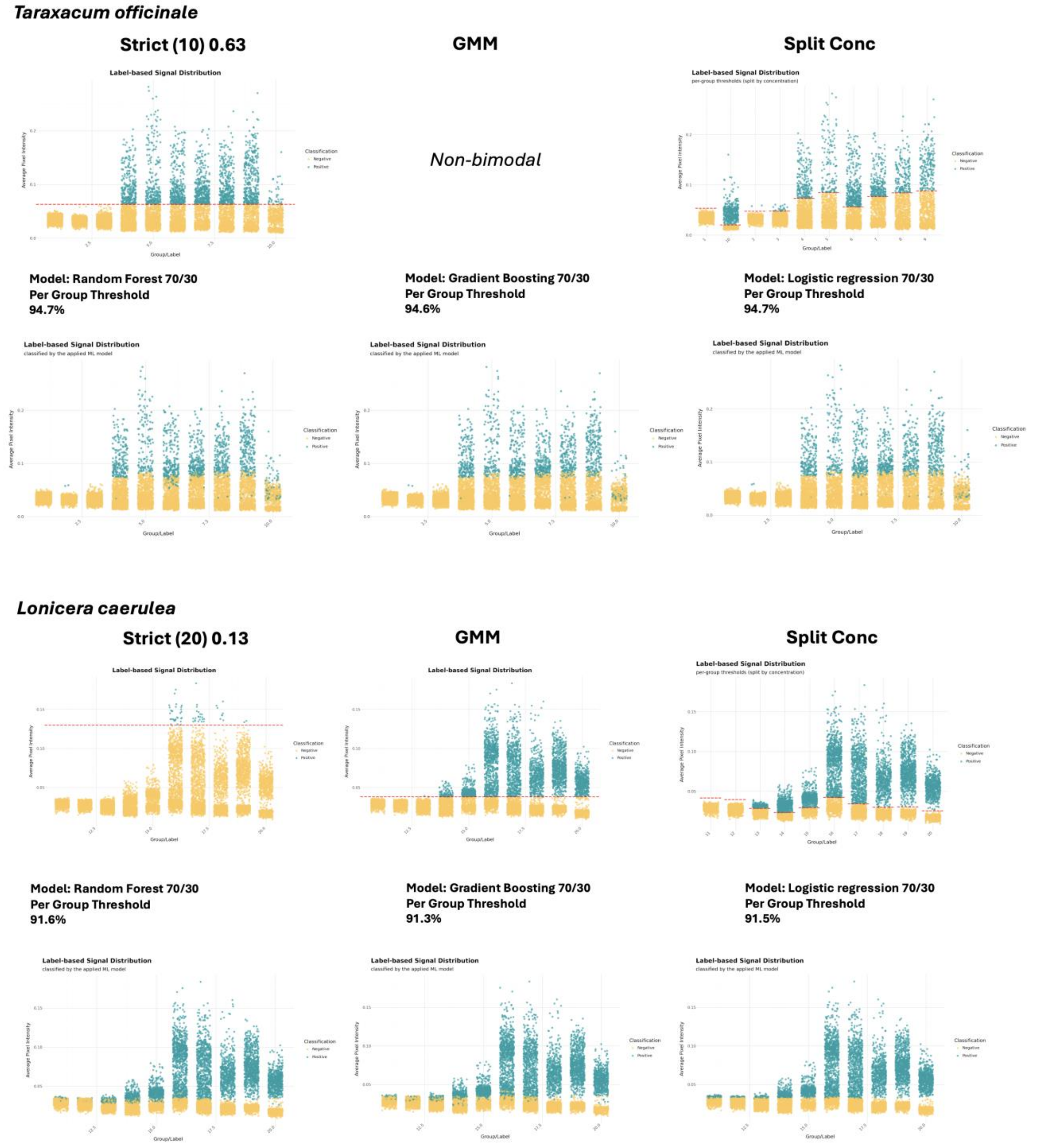


**Supplementary Figure S11.** Exploration of thresholding through Large Languange Models (LLMs) assistance in EasyFlow2 (A) *Taraxacum officinale* (B) *Lonicera caerulea*
